# Spotted fever group *rickettsiae* were the dominant pathogens in parasitic *Rhipicephalus microplus* in Yunnan Province, China

**DOI:** 10.1101/2023.09.24.559238

**Authors:** Zhang Lin, Miao GuangQing, Hou XueXia, Wang Peng, Guo Ying, Duan CunJuan, Yang XiaoNa, Hao Qin

## Abstract

**Objective:** Ticks as ectoparasites might carry many pathogens and transmitted to their hosts through blood feeding. Human might have the opportunity to be bite by ticks, therefore getting various tick-borne pathogens. Distribution of bacterium flora in ticks plays an essential role in mapping and preventing local tick-borne diseases. Our aim is to map the distribution of bacterium flora and identify the main tick-borne pathogens in dominant tick species in grazing areas of Yunnan province.

**Methods:** V3-V4 region of 16S rRNA amplifier sequencing was used to analyze and study the tick-borne pathogens in *Rh. Microplus*. The prevalence survey on *B.burgdorferi s.l*., *B. miyamotoi, E. chaffeensis, A. phagocytophilum, Coxiella burnetii* and Spotted fever group *rickettsiae* was carried out via PCR.

**Results:** The results showed 105 genus and 117 species were detected in 50 (24 in Yunpan, 26 in Menghun) ticks. Pathogen prevalence testing showed the ticks were positive for *B.burgdorferi* s.l.(9/50, 18%), Spotted fever group *rickettsiae* (35/50, 70%), and *A.phagocytophilum* (1/50, 2%).

**Conclusion:** In our analysis, Spotted fever group *rickettsiae* were the dominant pathogens in these areas. At the same time, we found that v3-v4 high-throughput sequencing of 16S rRNA gene is not sensitive to identify species for certain bacteria(such as *B. burgdorferi* s.l.). More accurate and comprehensive analysis would be necessary for detecting tick bacterium flora.

## INTRODUCTION

Ticks are obligate, blood-sucking ectoparasites that are commonly found all around the world. They are parasitic on amphibians, birds, reptiles and mammals. Some species can invade human beings and animals, and transmit many pathogens, including bacteria, viruses, and protozoa, which can cause many diseases and bring great harm to human health, animal husbandry production and wildlife ^1,2^. They are considered to be second only to mosquitoes as worldwide vectors of human disease ^3,4^.

Of the 117 described species in the Chinese tick fauna, 60 are known to transmit one or more diseases. Moreover, 38 of these species carry multiple pathogens, indicating the potentially vast role of these vectors in transmitting pathogens ^5^. We found that local dominant tick species was *Rhipicephalus Microplus* (*Rh. Microplus*) according to our surveillance in grazing areas in Yunnan province from 2017-2018 ^6^ and some reports ^7-9^. *Rh. Microplus* is an important tick species allover China, It can transmit several pathogens, such as *Anaplasma marginale, Babesia bigemina, Ba. bovis, Theileria equi*; *Ehrlichia chaffeensis* ; *tick-borne encephalitis virus (TBEV)* and *Coxiella burnetii* ^10^.

Yunnan is located in the southwest of China. Its geographical environment and climate are complex and diverse. According to the report, *Rh. Microplus* is widely distributed in Yunnan Province ^10,11^. *Rickettsia tsutsugamushi*, typhus, Q fever and *Bartonella* infection have been reported in Yunnan population ^12,13^. Yunpan is belonged to Pu’er city, which is famous for tea industy and covered by 67% forest area. Menghun in Xishuangbanna is located in the southern end of Yunnan Province. There are a large number of forest and pastoral areas suitable for tick growth. Xishuangbanna as a famous tourist area was attracted a large number of tourists every year. Therefore, the prevention and control of tick-borne diseases is particularly important. However, little research has been done on vector ticks and their pathogens in these areas. In order to fully understand the status of tick-borne pathogens and identify the main tick-borne pathogens in Yunpan and Menghun, and provide scientific basis for the prevention and control of tick-borne diseases, V3-V4 region of 16S rRNA amplifier sequencing was used to analyze and study the tick-borne pathogens in *Rh. Microplus* of these two areas.

## METHODS

### Tick collection and storage

More than 1000 parasitic ticks were collected from cattles in Yunpan and Menghun in March, 2018. These ticks were identified as *Boophilus microplus*. 24 *Rh. Microplus* ticks from Yunpan and 26 from Menghun were washed with 70% ethanol via gently shaking for 30 seconds, and then rinsed three times in sterile water to remove environmental contaminants ^14-16^ and frozen at -80°C until use.

### Genomic DNA extraction

Genomic DNA was extracted from 50 individual ticks using the QIAamp DNA Mini Kit according to the manufacturer’s protocols (QIAGEN, Shanghai, China). The extracted DNA was stored at -20°C until use.

### V3-V4 16S rRNA amplicon sequencing and data analysis

V3-V4 16S rRNA amplicon sequencing was performed by BGI Company, Shenzhen, China. The paired-end sequencing was performed on an Illumina Miseq platform (BGI, Shenzhen, China) based on a standard protocol from the manufacturer. Raw data were screened and assembled by QIIME^17^ and USEARCH (v7.0.1090)^3,18^ software. The UPARSE method was used to cluster the sequences into Operational Taxonomic Units (OTUs) at an identity threshold of 97%. Meanwhile, the RDP Classifier ^19^ was used to assign each OTU to a taxonomic level. Additional analysis such as rarefraction curves, Shannon index, and Good’s coverage, were carried out with QIIME.

### PCR amplification and sequence analysis

The prevalence survey on *B.burgdorferi s.l*., *B. miyamotoi, E. chaffeensis, A. phagocytophilum* and Spotted fever group *rickettsiae* was carried out via PCR, based on the genomic DNA extracted from individual ticks. All primer sequences and gene targets are listed in Table 1. The PCR products were separated using 1.5% agarose gel electrophoresis and visualized under UV light. PCR amplicons was sequenced by TsingKe Biological Technology Company, Beijing, China. The obtained sequences were queried to the NCBI BLASTn database to find the closest counterparts.

**Table 1.**
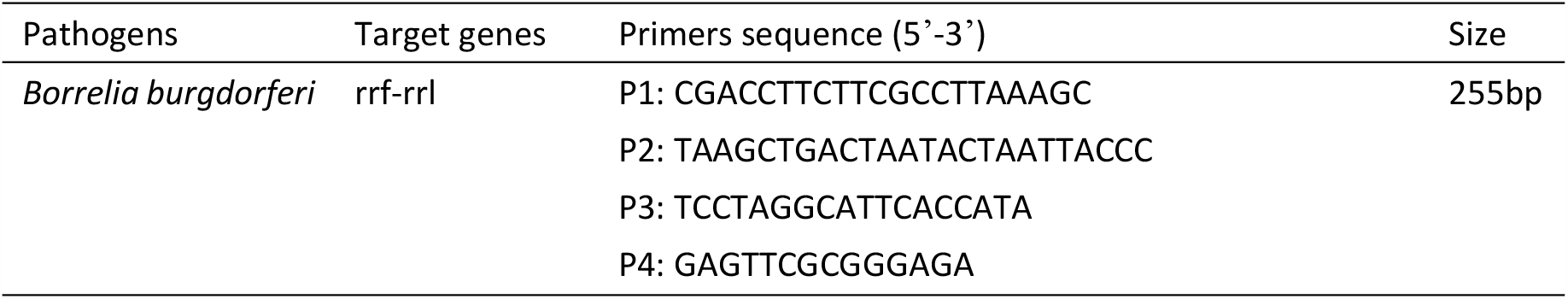
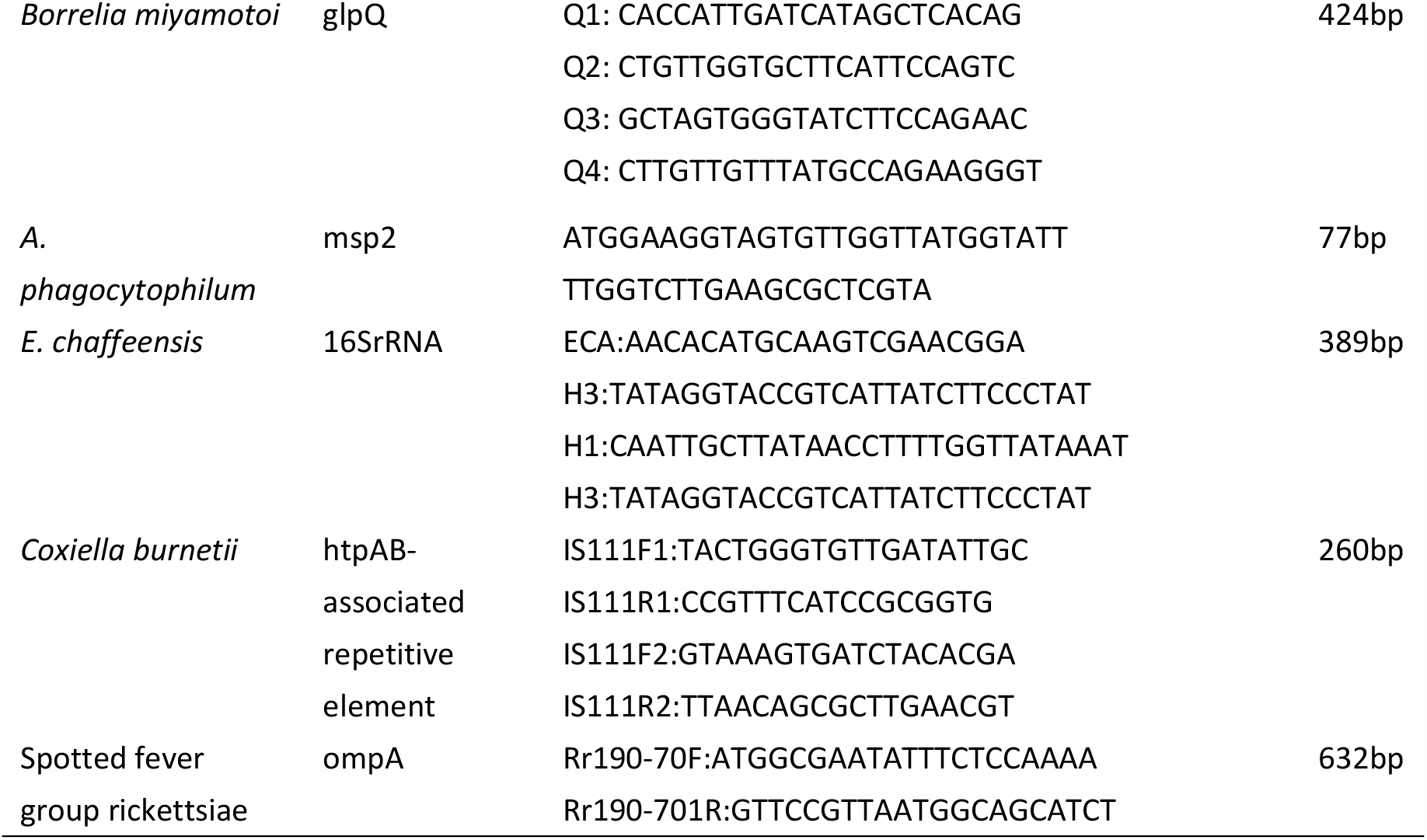
Target genes and primers of five tick-borne pathogens.

## RESULTS

### Bacterial community and diversity analysis

The V3-V4 region of 16S rRNA was sequenced in 50 tick samples. The results showed a total of 3,371,929 reads and 3,326,122 tags were obtained, the average length of reads was 67,438.58bp. There are 4844 OTUs produced by 50 tick copolymers.

At the phylum level, at least 19 phyla can be detected in 50 ticks, among which the four phyla with more contents are Proteobacteria, Bacteroidetes, actinobacteria and Firmicutes sequentially (Figure 1A). At the class level, the relative abundance of Gammaproteobacteria is the highest (Figure 1B). Spirochaetes was detected in three ticks (YN48, YN50 and YN72) at the phylum and class levels with a relative abundance of 0.005399%, 0.045022% and 0.023304% respectively.

**Figure 1.**
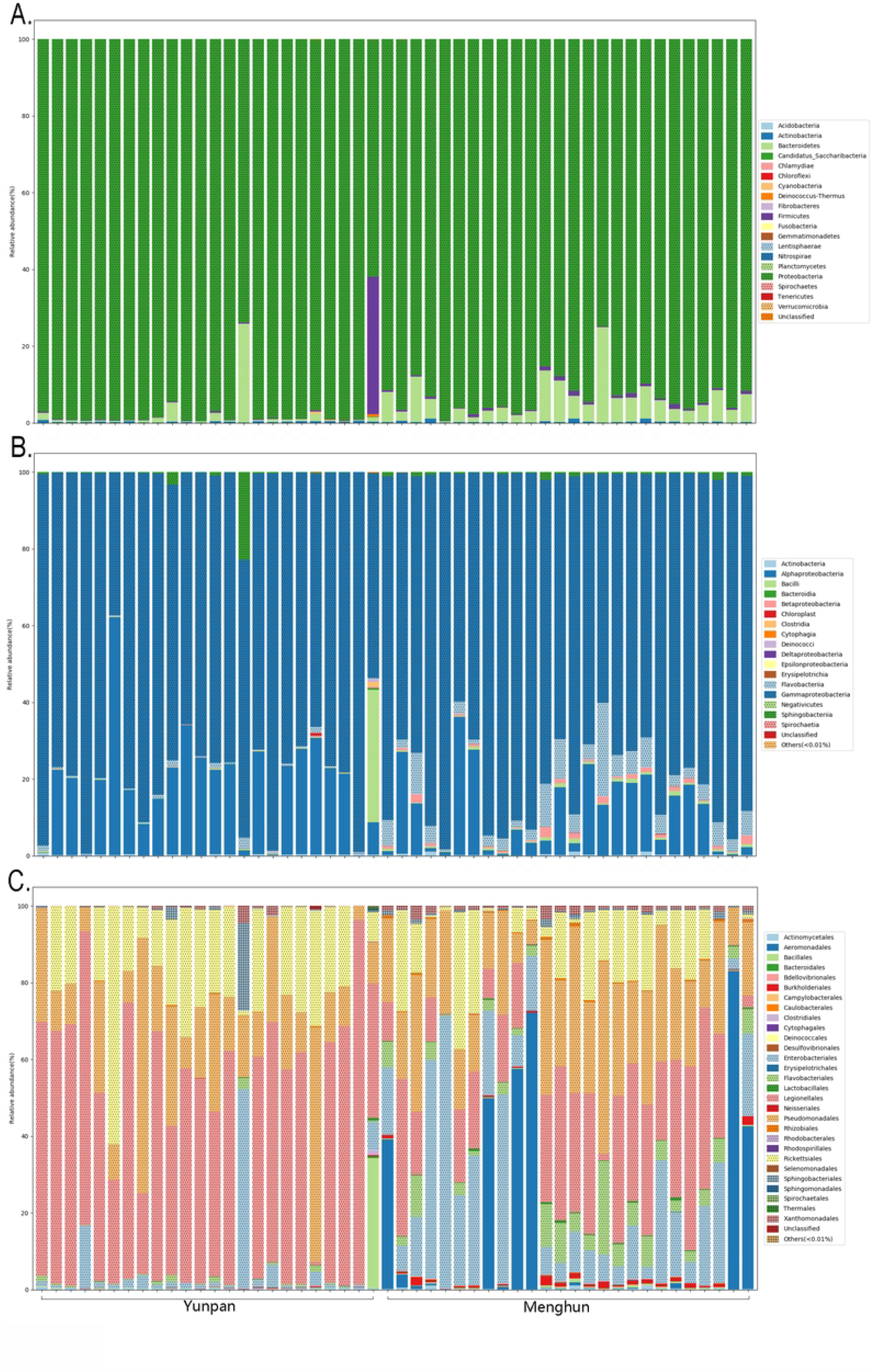
Bacterial community structure variation in 50 tick samples at A, the Phylum level; B, the Class level, and C, the Order level.

A total of 105 genus were detected in 50 ticks (Figure 2A). The micro-organisms of ticks have their own distribution characteristics at genus level, and the variation among individuals is large (Fig 2B). *Acinetobacter* was detected in all 50 tick samples, the relative abundance was 1.842026%∼44.86713%. *Aeromonas* was detected in 27 tick samples, the relative abundance was 38.91585% ∼ 82.95751%. *Anaplasma* was detected in 9 tick samples, among which the relative abundance of YN30 is the highest, reaching 51.09362%; the relative abundance of YN36 and YN46 were 3.699123% and 0.010648% respectively; the relative abundance of other six ticks was less than 0.01%. *Borrelia* was only detected in YN72 with relative abundance 0.023304%. *Coxiella* existed in all tick samples with relative abundance 0.011416% ∼ 95.07642%. *Ehrlichia* was detected in 27 ticks with relative abundance 0.001763% ∼ 35.84495%. *Enterococcus* was detected in 38 ticks, but the relative abundance was very low, only 0.001771% ∼ 0.280828%. *Pseudomonas* was detected in all tick samples, the relative abundance in YN32 and YN56 was 59.54905 and 24.9647. Rickettsia existed in 48 ticks, the relative abundance in YN35 was the highest (33.72437%). Staphylococcus existed in all ticks with the relative abundance 0.001893%∼0.208508%. Stenotrophomonas was detected in all 50 ticks, but the relative abundance in 13 ticks was higher(1.045084%∼4.415626%).

**Figure 2.**
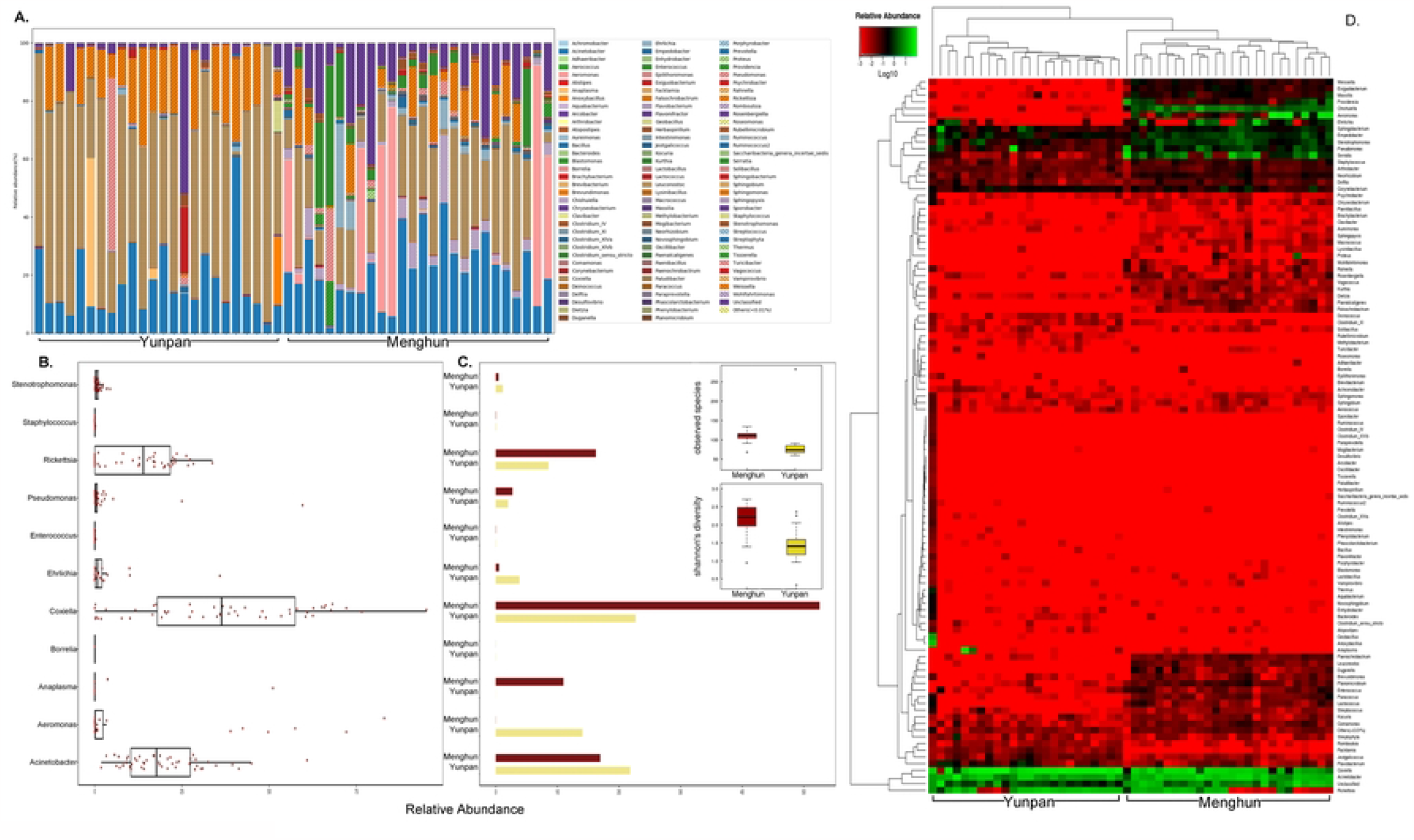
Bacterial community structure variation in 50 tick samples at Genus level. A, Relative abundance of barplot of all tick samples at Genus level; B, relative abundance of potential pathogenic bacteria in ticks and C, average abundance in Menghun County (maroon) and Yunpan County (yellow); D, heatmap of bacterial abundance in Genus level.

In addition, the results showed the high-probable pathogenic miro-organisms in all 50 samples were *Coxielia, Acinetobacter* and *Rickettsia* (Fig2B). The average abundance of the micro-organisms showed the similar pattern, while there were differences in two sample areas. It was higher that ticks in Menhun containing more *Coxilla* and *Rickettsia* than Yunpan, while with the *Acinetobacter* and *Aeromonas*, it showed the opposite way (Fig2C). More differences of tick microflora in Menghun and Yunpan happened when considering the relative abundance of *Weissella, Aeromonas* and *Ehrlichia*, etc. (Fig2D). At species level, A total of 117 species were detected in 50 ticks (Figure 3). Five *Acinetobacter* species including *Acinetobacter guillouiae, Acinetobacter johnsonii, Acinetobacter lwoffii, Acinetobacter ursingii* and *Acinetobacter variabilis* were detected in tick samples. *Aeromonas hydrophila* was detected in 27 ticks with the relative abundance 0.001814% ∼ 82.64443% and *Aeromonas salmonicida* was detected in 13 ticks with the relative abundance 0.001856%∼0.319997%.*Coxiella burnetii* accounts for a large proportion in the microbial composition of ticks. *Rickettsia heilongjiangensis* detected in 48 ticks was another important species. The proportions of *Anaplasma phagocytophilum* in YN30, YN36 and YN46 were 51.09362%, 3.699123% and 0.010648% respectively, and less than 0.01% in other 6 ticks. *Borrelia miyamotoi* was only detected in YN72 with the relative abundance 0.023304%. Four *Stenotrophomonas* species were detected in ticks, including *Stenotrophomonas chelatiphaga, Stenotrophomonas maltophilia, Stenotrophomonas rhizophila* and *Stenotrophomonas terrae*.

**Figure 3.**
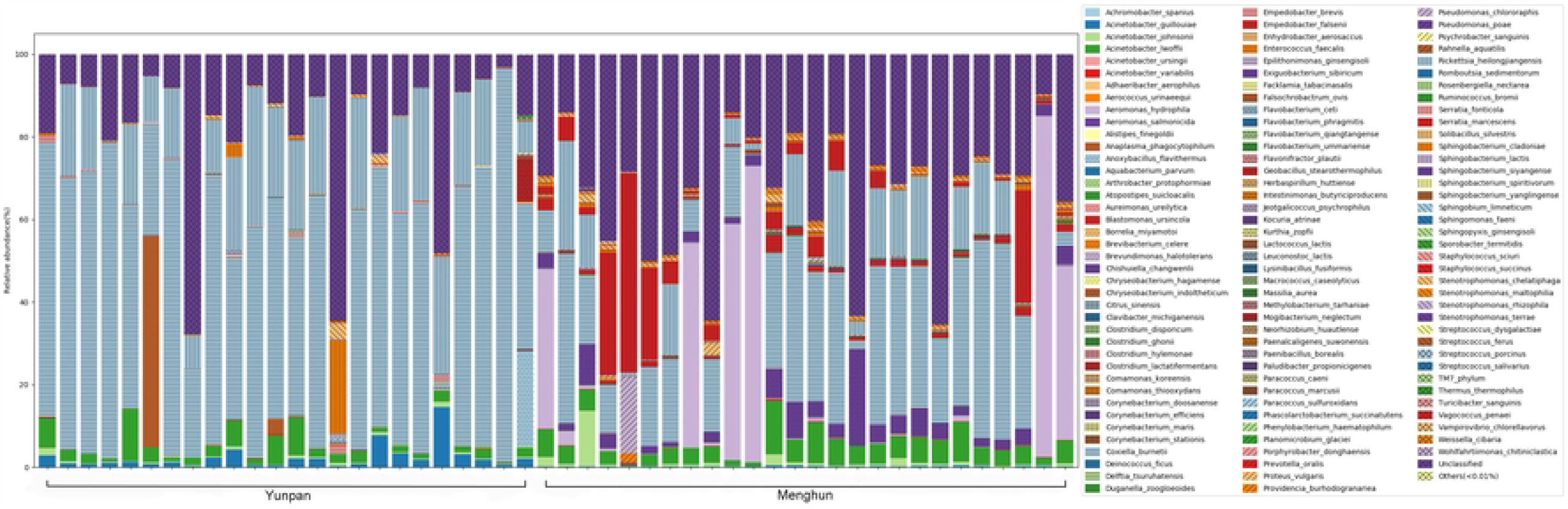
Bacterial community structure variation in 50 tick samples at the Species level

### Prevalence of tick-borne pathogen

In order to verify the prevalence of potential pathogens detected by high-throughout sequencing, we examined *B.burgdorferi s.l*., *B. miyamotoi, E. chaffeensis, A. phagocytophilum, Coxiella burnetii* and Spotted fever group *rickettsiae* in 50 ticks. The results showed 9 were positive for *B.burgdorferi s.l*.(18%, 7 in Yunpan and 2 in Menghun), 35 were positive for Spotted fever group *rickettsiae* (70%, 20 in Yunpan and 15 in Menghun), and 1 were positive for *A. phagocytophilum* (2%, in Yunpan). However, none of the *B.miyamotoi, Coxiella burnetii* or *E. chaffeensis* were detected in these 50 ticks by PCR (Table2), Although 16S rRNA amplifier sequencing results showed the existence. It might because that species detection by 16S rRNA amplifier sequencing may lead to misunderstanding for their accuracy or counts of reads. Therefore, verification using PCR or other methods are necessary.

**Table 2.** The results of 5 tick-borne pathogens in *Rh. Microplus*.

| Pathogens | No.of Ticks | Positives ( rates) | Yunpan |  | Menghun |  |
| --- | --- | --- | --- | --- | --- | --- |
|  |  |  | No. of Ticks | Positives (rates) | No. of Ticks | Positives (rates) |
| *B.b.s.l* | 50 | 9(18%) | 24 | 7(29%) | 26 | 2(8%) |
| *B.m* | 50 | 0 | 24 | 0 | 26 | 0 |
| *A.P.* | 50 | 1(2%) | 24 | 1(4%) | 26 | 0 |
| *E.C.* | 50 | 0 | 24 | 0 | 26 | 0 |
| *C. burnetii.* | 50 | 0 | 24 | 0 | 26 | 0 |
| *SFGR* | 50 | 35(70%) | 24 | 20(83%) | 26 | 15(58%) |
| *B.b.s.l&SFGR* | 50 | 7(14%) | 24 | 6(25%) | 26 | 1(4%) |
| *A.P. &SFGR* | 50 | 1(2%) | 24 | 1(4%) | 26 | 0 |

Additionally, we found that 7 ticks (14%) were co-infected with *B.burgdorferi s.l*. and Spotted fever group *rickettsiae*, and 1 tick (2%) was co-infected with *A. phagocytophilum* and Spotted fever group *rickettsiae*.

### The genospecies of B. burgdorferi s.l. and Spotted fever group rickettsiae

The results of 16S rRNA amplicon sequencing showed Spotted fever group *rickettsiae* were the dominant pathogens in ticks in either Yunpan or Menghun. PCR tests showed *B. burgdorferi s.l*. was an important pathogen in Yunpan. We determined *B. burgdorferi s.l*. and Spotted fever group *rickettsiae* in ticks. The 5S-23S rRNA intergenic region was sequenced for the positive ticks. The results showed that *B. burgdorferi s.l*. in 9 positive ticks were all belonged to *B.garinii. OmpA* gene was used to identify the Spotted fever group *rickettsiae*. The results showed that *candidatus rickettsia jiangxinensis* was the main subspecies in these areas.

## DISCUSSION

*Rh. Microplus* is the dominant tick species from cattle in Pu’er and Xishuangbanna, Yunnan province. Using 16S rRNA high-throughput sequencing method, the study of the microbial community and the pathogens in *Rh. Microplus* was carried out to provide scientific basis for the prevention and control of tick-borne diseases in local area. In this study 16S rRNA high-throughput sequencing was used to identify the full community of microbes and main pathogens within 50 parasitic *Rh. Microplus*, and also PCR methods were used to test the pathogens in these ticks. So we can have a comprehensive understanding of the microorganisms and pathogens in *Rh. Microplus* of this area.

It was found that the dominant microflora of parasitic *Rh. Microplus* in Menghun and Yunpan were *Proteobacteria, Bacteroidetes, actinobacteria* and *Firmicutes* sequentially. *Proteobacteria* is widely distributed in nature, including many pathogens, such as *E.coli, Salmonella, Vibrio, Helicobacter pylori* and other common species. *Bacteroidetes* plays an important role in carbohydrate fermentation, polysaccharide metabolism, bile acid and steroid metabolism, maintaining normal physiological function and microecological balance ^20^. It is of great significance to ticks.

Based on the analysis of population structure and diversity index, it was found that the species and relative abundance of micro-organisms carried by *Rh. Microplus* varied greatly, which was related to their feeding habits, males and females, and different growth stages. Studies have shown that there are some differences in the bacterial population structure of midgut contents between male and female *Rh. Microplus* in different stages ^21^.

In this study, conditional pathogens such as *acinetobacter, aeromonas* etc were detected with high relative abundance in some *Rh. Microplus. Acinetobacter* is one of the important opportunistic pathogenic bacteria to cause nosocomial infection. It can cause respiratory tract infection, septicemia, meningitis, endocarditis, urogenital tract infection, etc. *Aeromonas hydrophila* was also detected in 27 ticks with the relative abundance 0.001814%∼82.64443% . *Aeromonas hydrophila* is widely distributed in all kinds of water bodies in nature. It is the primary pathogen of many kinds of aquatic animals. Fish, frog and other cold-blooded animals are the natural hosts of the bacteria, which are the main sources of human infection. Patient carrier can also cause human to human transmission. If infected with *Aeromonas hydrophila*, the patients with low immune function are likely to have infection of endogenous blood, abdominal cavity, biliary tract, wound or urinary tract.

The results showed the genus of Spotted fever group *rickettsiae* were the dominant pathogens in Yunpan and Menghun. We detected Spotted fever group *rickettsiae* in 48 ticks by 16S rRNA sequencing with different relative abundances, and among which 35 were tested positive of *OmpA* gene by PCR. Previous study showed there were patients infected with *Rickettsia Siberia* in Yuxi area, Yunnan Province ^22^. Our study showed ticks of Pu’er and Xishuangbanna area were infected with *rickettsia jingxinensis*. The results suggested that there may be multiple subspecies of Spotted fever group *rickettsiae* in Yunnan Province. Of the genus of *coxiella* species, *Coxiella burnetii* was the pathogen of Q fever. Previous research showed patients with Q fever existed in some areas of Yunnan province ^12,13^. In this study, *Coxiella burnetii* was detected in 16S rRNA sequencing at species level, while it was not verified in PCR. It might because that results of 16 sRNA V3-V4 high-throughput sequencing at species level for *Coxiella burnetii* were not enough, therefore more verification would be needed. Our research shows that circulate of *Coxiella burnetii* between livestock and *Rh. Microplus* in these areas was at lower rate. Our results were also consistent with previous study that until now there is no any report of Q fever in Pu’er or Xishuangbanna area.

In this study no ticks were detected infection with *B. burgdorferi s.l*. by v3-v4 sequencing of 16S rRNA. However, we tested these 50 ticks with *B. burgdorferi s.l*. by PCR, the results showed that 9 ticks were positive. Sequencing of the 5S-23S rRNA intergenic region showed all the positive ticks were infected with *B. garinii*, which is one of the main pathogenic genotypes in China. This suggested v3-v4 sequencing of 16S rRNA is limited in the detection of species for certain pathogens.

In the 50 ticks, 7 ticks (14%) were co-infected with *B.burgdorferi s.l*. and Spotted fever group *rickettsiae*, and 1 tick(2%) was co-infected with *A. phagocytophilum* and Spotted fever group *rickettsiae*, indicating an increased risk of simultaneous human infection with these pathogens in Pu’er and Xishuangbanna area.

When focusing on the differences between Yunpan and Menghun, we noticed that the diversity of micro-organisms in Menghun was higher than Yunpan. However, considering the detection of pathogenic agents, we found the prevalence of *B. burgdorferi s.l*. and Spotted fever group *rickettsiae* in Yunpan was slightly higher than Menghun. Regional difference may exist as Yunpan is 400 km away from Menghun. This may need more investigation on dominant pathogenic agents in ticks to target ones properly and quickly in patients.

Through this study, we not only know the main microbial communities but also find the dominant pathogen of parasitic *Rh. Microplus* was Spotted fever group *rickettsiae* in Yunpan and Menghun using 16S rRNA high-throughput sequencing method. This provides basic data for local prevention and control of tick-borne disease. At the same time, we found that v3-v4 high-throughput sequencing of 16S rRNA gene is not sensitive to identify species for certain bacteria (such as *B. burgdorferi s.l*.).According to the report, while 16S rRNA sequencing is now widely used for microbial identification, this technique has been constrained by short read length of the most commonly used sequencing platform for microbial community, which often targeting only 1-3 variable regions in the 16S rRNA gene, such as V3-V5, V1-V3, or V4 alone. This constraint limits the taxonomic resolution ^23,24^. So if we want to get more accurate and comprehensive data, combination of sequencing multiple regions in the 16S rRNA gene may be necessary.

## CONFLICT OF INTEREST

The authors report no relationship that could be construed as conflict of interest.

## Notes

* This work was supported by Major Projects of the Thirteenth Five-Year Plan Special for Infectious Diseases (grant numbers 2018ZX10101002 and 2017ZX10303404-006-003).

The study is supported by the National Critical Project for Science and Technology on infectious Disease of P.R.China[2017ZX10303404-006-003]

### Competing Interest Statement

The authors have declared no competing interest.

